# Environmental and vector detection of African swine fever virus DNA in Vietnam: Evidence for potential transmission through wastewater and *Amblyomma javanense* ticks

**DOI:** 10.1101/2025.11.13.688377

**Authors:** Thi Ngan Mai, Duc Hieu Duong, Van Hieu Dong, Thi Huong Giang Tran, Thi Bich Phuong Cao, Thi My Le Huynh, Thanh Phong Bui, Van Kien Dao, Thi Hoa Nguyen, Thi Phuong Hoang, Tran Anh Dao Bui, Thi Lan Nguyen, Yasuko Yamazaki, Wataru Yamazaki

**Affiliations:** Department of Veterinary Microbiology and Infectious Diseases, Faculty of Veterinary Medicine, Vietnam National University of Agriculture, Hanoi Vietnam; Department of Veterinary Parasitology, Faculty of Veterinary Medicine, Vietnam National University of Agriculture, Hanoi Vietnam; Department of Veterinary Public Health, Faculty of Veterinary Medicine, Vietnam National University of Agriculture, Hanoi Vietnam; Hong Ha Nutrition Joint Stock Company, Ninhbinh Vietnam; Ceva Animal Health Vietnam, Hochiminh Vietnam; Key Laboratory of Veterinary Biotechnology, Faculty of Veterinary Medicine, Vietnam National University of Agriculture, Hanoi Vietnam; Department of Veterinary Pathology, Faculty of Veterinary Medicine, Vietnam National University of Agriculture, Hanoi Vietnam; Center for Southeast Asian Studies, Kyoto University, 46 Shimoadachi-cho, Yoshida, Sakyo-ku, Kyoto 606-8501, Japan

**Keywords:** African swine fever virus, wastewater, *Amblyomma javanense*, real-time PCR, Vietnam

## Abstract

African swine fever virus (ASFV) remains a major global threat to pig production, yet its environmental and vector reservoirs in Southeast Asia are poorly understood. This study provides the first evidence in Southeast Asia indicating that ASFV DNA can be detected in pig farm wastewater and that *Amblyomma javanense* ticks may act as potential mechanical carriers of the virus in natural settings. To elucidate part of the environmental dynamics and possible transmission routes of ASFV, we developed and applied a highly sensitive polyethylene glycol (PEG) precipitation-based real-time PCR method specifically for wastewater surveillance, as the viral load in aquatic environments is presumed to be substantially lower than that in arthropod vectors. A total of 40 tick samples and 93 wastewater samples were analyzed. ASFV DNA was identified in 3 of 40 ticks (7.5%) and 7 of 93 wastewater samples (7.5%). The developed method showed a 100-fold improvement in analytical sensitivity, lowering Ct values by 3–7 cycles and increasing the detection rate from 3.2% to 7.5% compared with the reference method. Diagnostic sensitivity and specificity for wastewater testing were 100.0% and 95.6%, respectively. Although infectious virus was not recovered from ASFV-positive wastewater samples, control experiments confirmed that PEG treatment did not interfere with viral infectivity. These findings highlight the potential role of environmental and tick-mediated transmission in ASF epidemiology and provide a practical tool for early warning and post-outbreak monitoring of ASFV in endemic regions.

## 1. Introduction

African swine fever (ASF) is a highly contagious viral disease of domestic pigs, causing severe economic losses and major impacts on the global swine industry (Sánchez-Vizcaíno et al. 2015; Galindo and Alonso 2017). Since its transboundary introduction into Georgia in 2007, the ASFV has spread across Europe, Russia, China, and Southeast Asia, resulting in devastating economic consequences and persistent endemicity (Rowlands et al. 2008; Sauter-Louis et al. 2021). Despite intensive control efforts, the absence of effective vaccines and limited diagnostic surveillance continue to hinder eradication (Blome et al. 2020; Dixon et al. 2020).

In Africa, warthogs and soft ticks of the genus *Ornithodoros* play crucial roles as long-term ASFV reservoirs, enabling viral maintenance in sylvatic cycles (Jori and Bastos 2009; Penrith et al. 2019). However, in Southeast Asia, the ecological reservoirs of ASFV remain poorly characterized, as few studies have investigated the potential roles of local tick species or environmental contamination in virus persistence. In particular, the possible involvement of hard ticks such as *Amblyomma javanense* and the role of contaminated wastewater in ASFV maintenance have not yet been confirmed in the region. Environmental transmission routes, particularly through wastewater and effluents from infected farms, have been hypothesized but remain unverified (Le et al. 2019; Nguyen et al. 2021).

Recent epidemiological analyses in India suggested that illegal disposal of infected pig carcasses into river systems could contribute to ASFV dissemination (Bora et al. 2020; Buragohain et al. 2023). However, these findings were based solely on indirect epidemiological inference, and to date, no study has provided direct molecular evidence of ASFV DNA in farm effluents or environmental waters in Southeast Asia. Similarly, the potential mechanical role of *Amblyomma javanense* ticks in ASFV transmission has not been explored.

Given the increasing frequency of ASF outbreaks and the limited understanding of its environmental persistence, integrating vector and environmental surveillance represents a crucial but underexplored component of ASFV epidemiology. Therefore, this study aimed to elucidate part of the environmental dynamics and potential transmission routes of ASFV in Vietnam by simultaneously investigating the presence of ASFV in *Amblyomma javanense* ticks and pig farm wastewater. In doing so, we also developed and validated a highly sensitive polyethylene glycol (PEG)-based real-time PCR method for detecting ASFV in complex environmental matrices (Mai TN et al. 2025), providing a new tool for integrated vector and environmental surveillance in endemic regions.

## 2. Material and method

### 2.1. Ethics approval

All experiments were approved by the Animal Research Ethics Committee (CARE) of the Faculty of Veterinary Medicine, Vietnam National University of Agriculture (Approval No. CARE-2022/05). All procedures were conducted in accordance with the ARRIVE guidelines and complied with the WOAH (World Organisation for Animal Health) standards for animal welfare.

### 2.2. Pig farm wastewater samples

#### Sample collection

Wastewater samples were collected from 16 commercial pig farms (9 finishing and 7 sow farms) across northern Vietnam between March 2023 and June 2025 for the detection of ASFV. Sampling followed the guidelines of the World Organisation for Animal Health (WOAH) and the European Union Reference Laboratory for ASF (EURL-ASF). In finishing farms, samples were obtained from pens with suspected cases and adjacent pens, specifically from drainage points or wastewater outlets. In sow farms, samples were collected directly from centralized wastewater storage tanks. Each sample (50–200 ml) was collected in sterile plastic tubes, kept at 4 °C during transportation, and processed within 24 h at the Key Laboratory for Veterinary Biotechnology, Faculty of Veterinary Medicine, Vietnam National University of Agriculture (VNUA).

#### DNA extraction

##### Reference method

To remove coarse debris and potential PCR inhibitors, raw wastewater samples were centrifuged at 2,000 *g* for 10 min. A 150 μl aliquot of the resulting supernatant was subjected to DNA extraction using the DNeasy Blood & Tissue Kit (Qiagen, Maryland, USA) and eluted in 50 μl of buffer, following the manufacturer’s protocol.

##### Developed PEG-based method

A PEG-based virus concentration method, adapted from Mai et al. (2025), was optimized for ASFV detection in wastewater. Briefly, 15 ml of raw wastewater was centrifuged at 2,000 *g* for 10 min to remove debris. Then, 13 ml of clarified supernatant was mixed with 5.2 ml of PEG–NaCl solution (final PEG concentration ≈ 40%), vortexed, and centrifuged at 8,000 *g* for 20 min. The pellet was resuspended in 150 μl of PBS, followed by a secondary centrifugation at 9,000 *g* for 3.5 min. The final supernatant (150 μl) was subjected to DNA extraction using the DNeasy Blood & Tissue Kit (Qiagen), as above.

#### Real-time PCR assay

Detection of ASFV DNA extracted from wastewater samples was performed using real-time PCR on the LineGene Mini S real-time PCR system (Bioer Technology, Hangzhou, China). The assay employed a set of primers and probe targeting the ASFV p72 gene, as described by Tignon et al. (2011), including a forward primer (5’-TGC TCA TGG TAT CAA TCT TAT CG-3’), a reverse primer (5’-CCA CTG GGT TGG TAT TCC TC-3’), and a fluorescently labeled probe (5’-FAM-TTC CAT CAA AGT TCT GCA GCT CTT-TAMRA-3’). Each reaction was prepared in a final volume of 25 µl, containing: 12.5 µl of 2× probe qPCR mix (Takara Bio Inc., Otsu, Japan), 0.1 µl each of the forward and reverse primers (100 µM; Hokkaido System Science, Sapporo, Japan), 0.05 µl of probe (100 µM, Hokkaido System Science), 7.25 µl of nuclease-free water (Takara Bio), and 5 µl of DNA template extracted from the concentrated wastewater sample. Thermal cycling conditions were as follows: initial denaturation at 95 °C for 30 seconds, followed by 45 cycles of denaturation at 95 °C for 10 seconds and annealing/extension at 60 °C for 60 seconds. The cycle threshold (Ct) values were determined automatically by the instrument software, with a cut-off value of 37.00 used to define positive ASFV detection. Each run included a positive DNA control, a negative extraction control, and a no-template control to ensure assay integrity.

#### Determination of the limit of detection

The ASFV strain ASF/HY01/2019 (GenBank Accession No. MK554698; genotype II) was propagated in porcine alveolar macrophages and stored at –80 °C with a titer of 10^6,5^ TCID_50_. All ASFV manipulations were conducted at the Key Laboratory of Veterinary Biotechnology, Vietnam National University of Agriculture. Prior to use, the viral stock was thawed and serially diluted 10-fold in phosphate-buffered saline (PBS), with all dilutions kept on ice during handling. Concurrently, raw wastewater samples previously stored at –80 °C were thawed at room temperature and centrifuged at 2,000 *g* for 10 minutes to remove solids and debris. The clarified supernatant was divided into sterile 50 ml conical tubes (15 ml per tube). Each ASFV dilution, ranging from undiluted to 10⁻⁵, was added to a separate aliquot and vortexed thoroughly to ensure even distribution of the virus.

This dilution series was used to assess viral recovery, concentration efficiency, and the analytical sensitivity of the detection protocol, including DNA extraction and real-time PCR. DNA was extracted from each spiked sample and tested in duplicate. A sample was considered positive when both replicates produced amplification signals with Ct values below the threshold. The limit of detection (LOD) was defined as the lowest virus concentration at which consistent amplification was observed in both replicates. The limit of detection (LOD) was defined as the lowest dilution consistently detected in both replicates (Ct < 37.0).

### 2.3. Tick samples

#### Sample collection

In this study, a total of 40 ticks were collected from Sunda Pangolins *(Manis javanica)* rescued at the Rescue Center within Cuc Phuong National Park (Ninh Binh Province, Vietnam) between Apri and August, 2024. The animals were confiscated from illegal wildlife trade and kept under quarantine before release. Tick collection was performed manually during the quarantine period at the rescue station using sterile forceps to remove ticks from infestation sites to avoid damaging mouthparts. Collected ticks were first morphologically examined under a stereomicroscope and identified based on standard taxonomic keys described by Guglielmone et al. (2014) to confirm species identity. After morphological identification, ticks were preserved in sealed plastic tubes containing RNA preservation solution or kept frozen in liquid nitrogen depending on field conditions. All samples were properly labeled and transported under refrigerated conditions to the laboratory for further analysis.

#### DNA extraction from ticks using liquid nitrogen and CTAB buffer

Genomic DNA (gDNA) was extracted using a modified cetyltrimethylammonium bromide (CTAB) method (Doyle and Doyle 1987). This approach is particularly suitable for arthropod tissues, as mechanical grinding under liquid nitrogen efficiently disrupts the chitinous exoskeleton, while CTAB effectively removes complex polysaccharides and other PCR inhibitors commonly present in tick samples. Briefly, individual ticks were flash-frozen in liquid nitrogen and homogenized with 1 ml of 1% CTAB buffer. The homogenate was incubated at 65 °C for 60 min following the addition of 120 µl of 10% SDS and 10 µl of β-mercaptoethanol. After centrifugation (13,700 *g*, 10 min, 4 °C), the supernatant was extracted once with PCI (phenol:chloroform:isoamyl alcohol, 25:24:1) and twice with CI (chloroform:isoamyl alcohol, 24:1). DNA was precipitated with cold isopropanol (−20 °C, 15 min) and pelleted by centrifugation (10,000 *g*, 7 min, 4 °C). The pellet was washed two to three times with cold 70% ethanol, air-dried, and resuspended in 50 µl of TE buffer (pH 8.0). DNA samples were stored at -20^0^C until further use.

#### Real-time PCR assay

Detection of ASFV DNA extracted from tick samples was performed using real-time PCR on the LineGene Mini S real-time PCR system (Bioer Technology, Hangzhou, China) as described above.

### 2.4. Statistical analysis

The diagnostic sensitivity, specificity, positive predictive value (PPV), and negative predictive value (NPV) were estimated. Differences in paired proportions were assessed using McNemar’s test, performed in R software (version 4.5.1). A p-value of less than 0.05 was considered statistically significant. Confidence intervals (95% CI) were calculated using the Wilson score method.

## 3. Results

### 3.1. The limit of detection for ASFV detection methods in wastewater samples

As shown in Table 1, spiking experiments demonstrated that the developed real-time PCR method exhibited approximately 100-fold higher analytical sensitivity than the reference method for ASFV detection in wastewater samples. The developed assay consistently detected viral DNA at dilutions up to 10⁻⁴, while the reference method’s detection limit was at 10⁻² dilutions. These results highlight the improved analytical sensitivity of the developed method for identifying low concentrations of ASFV in complex wastewater matrices.

**Table 1.** Limit of detection (LOD) of the developed and reference methods for ASFV detection in spiked ASFV-negative pig farm wastewater samples.

| Assay | Dilutions of pooled ASFV-spiked wastewater samples |  |  |  |  |  |
| --- | --- | --- | --- | --- | --- | --- |
|  | 10<sup>0</sup> | 10<sup>-1</sup> | 10<sup>-2</sup> | 10<sup>-3</sup> | 10<sup>-4</sup> | 10<sup>-5</sup> |
| Real-time PCR (Ct) |  |  |  |  |  |  |
| Developed method | 24.54 | 28.28 | 31.91 | 33.75 | 37.00 | No. Ct |
| Reference method | 28.59 | 31.96 | 35.61 | No. Ct | No. Ct | No Ct |

Ct, Threshold cycle. No. Ct, no threshold cycle values detected by real-time PCR. All results were consistent in duplicate testing, with no discordant outcomes.

### 3.2. Surveillance of African swine fever virus in farm wastewater samples

A total of 93 wastewater samples were analyzed to evaluate the diagnostic performance of the developed method in comparison with the reference method. As shown in Table 2, all three samples identified as positive by the reference method were also detected as positive by the developed method. Notably, the Ct values obtained with the developed method were consistently lower than those from the reference method, with differences ranging from 3.05 to 7.09 cycles. According to the principle of real-time PCR, each cycle reduction corresponds to an approximate two-fold increase in detectable viral genetic material, indicating an increase in viral load of approximately 8- to 128-fold (2³ to 2⁷). These findings suggest that the developed method offers improved sensitivity in detecting ASFV, particularly in samples with low viral loads.

**Table 2.** Diagnostic performance of real-time PCR using the developed and reference DNA extraction methods for wastewater samples.

|  | Reference positive<br>(*n* = 3) | Reference negative<br>(*n* = 90) |
| --- | --- | --- |
| Developed positive (*n* = 7) | 3 | 4 |
| Developed negative (*n* = 86) | 0 | 86 |

Diagnostic sensitivity, 100% (3/3; 95% CI: 29.2–100.0%). Diagnostic specificity, 95.6% (86/90; 95% CI: 89.2–98.8%). PPV, 42.9% (95% CI: 9.9-81.6%). NPV, 100.0% (95% CI: 95.8–100.0%). Developed method: improved DNA extraction combined with real-time PCR; Reference method: standard DNA extraction followed by real-time PCR. Results were consistent in duplicate testing.

In addition, four samples that tested negative by the reference method were newly identified as positive by the developed assay, resulting in a total of seven positive samples (7.5%) detected by the developed method, compared to only three positive samples (3.2%) by the reference method. All four samples tested negative using the reference method but produced clear amplification curves with Ct values below the cutoff in the developed assay. The diagnostic sensitivity and specificity of the developed real-time PCR method were 100.0% (95% confidence interval [CI]: 29.2–100.0%) and 95.6% (95% CI: 89.2–98.8%), respectively. The positive predictive value (PPV) was 42.9% (95% CI: 9.9–81.6%), while the negative predictive value (NPV) was 100.0% (95% CI: 95.8–100.0%). The increase in detection rate achieved by the developed method was statistically significant (p = 0.046) by McNemar’s test, indicating improved sensitivity and potential for ASFV detection in wastewater samples compared to the reference protocol.

To determine whether the ASFV DNA detected represented infectious virus or merely viral genetic material, all positive wastewater samples were subjected to virus isolation attempts in the PAM cell culture. However, none of the culture attempts yielded positive results, likely due to the low viral loads present in these samples (data not shown). To validate the virus isolation process and exclude inhibitory effects of PEG used during sample processing, control experiments were performed by inoculating cells with ASFV mixed with PEG. The positive control wells exhibited clear cytopathic effects, confirming that PEG did not interfere with viral infectivity or culture conditions (data not shown). These findings suggest that while the developed method is highly sensitive in detecting ASFV DNA in wastewater, the infectious virus may be present at levels below the threshold for successful isolation.

Notably, the four additional positive samples identified exclusively by the developed method provide important epidemiological insights. Two samples were collected from a sow farm that had experienced an ASF outbreak four months prior. Although the facility had undergone thorough cleaning, disinfection, and a fallow period following the outbreak, ASFV DNA was still detectable in wastewater samples, suggesting potential long-term environmental persistence of viral genetic material. The third sample was obtained from the wastewater of a 600-head finishing pig farm with a history of recurrent ASF outbreaks, further highlighting the potential of the developed assay to detect residual or low-level viral contamination in high-risk environments. Following the positive result by the developed method, the farm implemented enhanced disinfection and biosecurity measures. Notably, no further ASF outbreaks have occurred to date, suggesting that early detection of viral traces via wastewater surveillance can support timely interventions and effective outbreak prevention. This case demonstrates the practical value of wastewater-based surveillance for ASFV, providing a non-invasive, proactive tool to monitor environmental contamination and guide targeted disease control efforts, especially in farms with previous infections or heightened transmission risk.

The remaining sample was obtained from a 600-pig farm located in Bac Giang province that had previously reported ASF outbreaks. Among the 86 concordant negative samples, 13 samples recorded Ct values between >37 and <40 with the developed method, falling within the assay’s suspicious range. These borderline signals may indicate trace levels of viral DNA near the detection limit, underscoring the higher analytical sensitivity of the developed method.

### 3.3. Detection of African swine fever virus in ticks

A total of 40 tick samples were identified as *Amblyomma javanense*, then were tested for the presence of ASFV DNA using real-time PCR. Among these, three samples (7.5%) tested positive, indicating the presence of ASFV genetic material in the tick population associated with confiscated Sunda pangolins at the Cuc Phuong Rescue Center. The three positive samples were collected from individual ticks originating from different host animals, and Ct values ranged from 34.66 to 36.80 (Table 3), suggesting low to moderate viral loads.

**Table 3.** Summary of ASFV-positive tick samples detected by real-time PCR.

| **Sample ID** | **Host ID** | **Ct value** | **Tick stage** | **Collection date** |
| --- | --- | --- | --- | --- |
| PWA221901.1 | Pangolin-07 | 34.66 | Adult | April, 2024 |
| PWA221906.13 | Pangolin-11 | 36.80 | Adult | April, 2024 |
| PWA221906.14 | Pangolin-14 | 35.68 | Adult | April, 2024 |

Ct, Threshold cycle. All results were consistent in duplicate testing.

Notably, only one of the three positive samples ID PWA221901.1 showed a weak positive signal (Ct = 34.66) after two rounds of PCR amplification targeting the P72 gene. Although the PCR product was subsequently subjected to cloning and sequencing attempts, no sequencing product was obtained, likely due to the low viral load and insufficient amplicon quantity. This is consistent with the high Ct value observed, which typically reflects a low concentration of viral DNA in the sample. These findings provide the first molecular evidence of ASFV DNA in *Amblyomma javanense* ticks from wildlife rescue settings in Vietnam. While the detected viral loads were low, these results suggest that ticks associated with wildlife species may serve as potential mechanical carriers or environmental reservoirs for ASFV in natural ecosystems.

## 4. Discussion

ASFV continues to pose a persistent challenge for global pig production, not only through direct transmission among domestic herds but also through poorly understood environmental and vector reservoirs. This study provides the first molecular evidence from Southeast Asia that ASFV DNA can be detected in both pig farm wastewater and *Amblyomma javanense* ticks in Vietnam, suggesting that these compartments may contribute to viral persistence and potential indirect transmission. By addressing this knowledge gap, our investigation aimed to elucidate part of the environmental dynamics and transmission pathways of ASFV, thereby expanding the current understanding of its ecology beyond the host population. This is, to our knowledge, the first integrated investigation of ASFV presence in both environmental and tick samples in the region, underscoring the importance of combining vector and environmental surveillance with conventional diagnostic approaches for more comprehensive ASF control.

In the spike experiments, the developed real-time PCR method demonstrated approximately 100-fold greater sensitivity compared to the reference assay for detecting ASFV in wastewater samples. This marked improvement is primarily attributed to the optimized sample preparation protocol, which incorporates a preliminary centrifugation step to remove solid debris and potential PCR inhibitors, followed by virus concentration using PEG precipitation combined with centrifugation. Such optimization enables efficient nucleic acid recovery even from complex matrices, consistent with prior studies showing that concentration and purification steps significantly improve viral detection sensitivity (Kwon et al. 2024; Monteiro et al. 2024). Kwon et al. (2024) reported significant improvements in ASFV DNA recovery from environmental samples containing organic inhibitors by employing refined pretreatment methods, including centrifugation, filtration, and magnetic-bead purification. Likewise, Monteiro et al. (2024) demonstrated effective wastewater-based surveillance for viral pathogens such as dengue and chikungunya, incorporating a virus concentration step using hollow-fiber filtration that proved essential for enhancing detection sensitivity and reliable monitoring of viral circulation in community wastewater. Overall, these methodological enhancements, including effective sample pretreatment and optimized virus concentration techniques, are crucial for reliable molecular detection of ASFV under field conditions and support early outbreak surveillance through environmental monitoring.

The developed real-time PCR method exhibited superior sensitivity in detecting ASFV in wastewater samples, identifying seven positive samples compared to three detected by the reference assay. Diagnostic sensitivity reached 100%, with a specificity of 95.6%, consistent with previous findings emphasizing the importance of optimized viral concentration and nucleic acid extraction in complex environmental matrices (La Rosa et al. 2020). Among the 86 samples negative by both methods, 13 showed borderline Ct values between 37 and 40 with the developed assay, indicating low-level viral presence near the detection limit. Notably, three of these samples were collected from farms where ASF outbreaks occurred within days after sampling, emphasizing the need for cautious and follow-up testing in such epidemiological contexts.

These findings align with recent advances in wastewater-based surveillance, which has been increasingly recognized as an effective approach for early detection and monitoring of viral circulation in communities, particularly in low-prevalence or subclinical scenarios (Ahmed et al. 2020; Medema et al. 2020). The higher detection rate observed here suggests that the developed method enhances surveillance sensitivity by capturing low-level viral shedding missed by conventional assays. Notably, four additional positive wastewater samples identified exclusively by the developed method originated from distinct epidemiological contexts: two from a sow farm sampled four months after an ASF outbreak despite thorough cleaning and disinfection; one from a finishing pig farm with recurrent ASF outbreaks where enhanced biosecurity measures were subsequently implemented; and one from a region in Bac Giang with recent sporadic cases. These findings illustrate the assay’s applicability across diverse field settings and its ability to detect residual viral DNA in environments with varying contamination histories. The increased sensitivity of the developed method was evident in three concordant positive field samples, where Ct values were reduced by 3.05 to 7.09 cycles compared to the reference assay, corresponding to an approximately 8- to 128-fold increase in detectable viral load. These results highlight the effectiveness of the optimized sample preparation steps specifically, debris removal by centrifugation and viral concentration via PEG precipitation in enhancing ASFV detection from wastewater matrices.

While detection of viral DNA in wastewater does not confirm viral infectivity or transmission risk, it highlights potential environmental persistence consistent with ASFV’s known stability in organic-rich matrices such as manure and soil, especially under favorable conditions (Gallardo et al. 2019; Mazur-Panasiuk et al. 2019). ASFV is known to survive for extended periods in environmental matrices ((Davies et al. 2017; Gu et al. 2025). In many pig farms, drinking water is often unchlorinated or poorly disinfected to reduce operational costs. Consequently, ASFV present in farm wastewater may contaminate drinking-water sources of downstream farms, posing a serious risk for secondary spread of infection (Mai NT et al. 2022; U.S. Department of Homeland Security 2025). This underscores the utility of sensitive molecular assays for both outbreak detection and post-eradication environmental monitoring.

Although hard ticks of the genus *Amblyomma* have not been confirmed as biological vectors of ASFV, unlike soft ticks of the genus *Ornithodoros* which are well-established in the sylvatic transmission cycle of the virus in Africa (Penrith 2009; Brown and Bevins 2018) the detection of ASFV DNA in *Amblyomma javanense* ticks collected from rescued *Manis javanica* pangolins in this study provides meaningful epidemiological insights. This finding indicates that ASFV may be present not only in domestic pig populations but also within wildlife-associated ecosystems. It expands the current understanding of ASFV ecology in Vietnam, indicating that the virus may persist or circulate in natural environments beyond farm settings.

Public information about the Rescue Center in Cuc Phuong National Park indicates that the facility primarily houses confiscated and rescued wildlife and operates strict quarantine protocols for incoming animals (Cuc Phuong National Park 2024). We were not able to identify any public record suggesting the presence of domestic pigs or wild boars within the center. Accordingly, while the detection of ASFV DNA in *Amblyomma javanense* ticks collected from rescued pangolins may reflect mechanical carriage acquired under natural conditions, we cannot exclude the possibility that the ticks or their hosts had prior contact with infected suids before rescue. Further verification of capture histories and quarantine records, along with targeted ecological studies, will be valuable to clarify whether *Amblyomma javanense* can act as a natural mechanical carrier of ASFV.

Supporting this interpretation, an ASF outbreak in wild boar was repoted between late November and December 2024, during which 29 wild boars were found dead in Pù Mát National Park, Vietnam ; subsequent laboratory testing confirmed ASFV infection (Hung 2024). This event highlights that wildlife populations in Vietnam can be directly affected by ASFV, reinforcing the relevance of wildlife–tick surveillance within a One Health framework. Herein, *Amblyomma javanense* ticks may serve as an environmental indicator or an “epidemiological footprint” of ASFV exposure among wildlife and their habitats. Therefore, monitoring ASFV in wildlife-associated ticks represents a practical surveillance strategy, enabling early detection of viral circulation at the wildlife–environment interface before potential spillover into domestic swine populations occurs.

This study has several limitations. First, the relatively small sample size of ticks (*n*=40) and wastewater samples (*n*=93) may limit the generalizability of the findings across broader geographic areas or diverse farming systems. Second, the detection of ASFV DNA does not necessarily indicate the presence of infectious virus, as viral isolation was unsuccessful in all positive wastewater samples-likely due to low viral loads. Third, sequencing attempts failed due to insufficient amplicon yield, preventing confirmation of ASFV genotypes and phylogenetic relationships. Fourth, because the *Amblyomma javanense* ticks were collected from rescued pangolins at a wildlife rehabilitation facility, the possibility of prior exposure of the hosts or ticks to infected suids before rescue cannot be fully excluded. Future studies with larger sample sizes, virus isolation, and genomic characterization are warranted to better elucidate the role of ticks and wastewater in ASFV epidemiology in Vietnam.

Overall, this study demonstrates the feasibility of ASFV detection through integrated environmental and vector surveillance. The developed PEG-based real-time PCR assay offers enhanced analytical sensitivity for low-titer samples, enabling early detection of ASFV DNA in complex matrices. By providing the first molecular evidence of ASFV DNA in both wastewater and *Amblyomma javanense* ticks in Southeast Asia, this study contributes new insights into the environmental dynamics and potential transmission routes of ASFV. These findings emphasize the importance of adopting an integrated “One Health” approach that links animal, environmental, and wildlife health, thereby strengthening ASF control and early-warning systems in endemic regions.

## 5. Conclusion

This study provides the first integrated molecular evidence in Southeast Asia of ASFV DNA in both pig farm wastewater and *Amblyomma javanense* ticks in Vietnam, revealing potential environmental and vector pathways for virus persistence. The newly developed PEG-based real-time PCR assay demonstrated a 100-fold improvement in detection sensitivity compared with the reference method, enabling the identification of low-level ASFV contamination in post-outbreak farms and complex environmental matrices. These results not only clarify part of the environmental dynamics of ASFV but also establish a foundation for future integrated surveillance in endemic areas. Integrating wastewater and tick-based monitoring into national surveillance frameworks, guided by a One Health perspective, could substantially strengthen biosecurity measures and early warning systems for transboundary animal diseases in the region.

## Data availability

The data that support the findings of this study are available from the correspondence author upon reasonable request.

## Acknowledgements

We would like to thank the field veterinarians for their valuable support in field sample collection.

## Author contributions

MTN designed the study, performed the laboratory testing, statistical analysis, and drafted the manuscript. DDH collected tick samples and drafted the manuscript. DVH and TTHG collected wastewater samples and performed the laboratory testing. CTBP and HTP participated in the the laboratory testing. BTP and DVK collected wastewater samples. HTML, BTAD and NTL designed the study. YY and WY participated in the design of the study, drafted and reviewed the manuscript. WY received funding from JSPS KAKENHI. All authors have reviewed and approved of the submitted version.

